# Choosing safety or success: Fall-avoidance preference limits goal achievement during whole-body movements

**DOI:** 10.1101/2025.07.02.662696

**Authors:** Taisei Fujimura, Shota Hagio, Motoki Kouzaki

**Affiliations:** Laboratory of Neurophysiology, Graduate School of Human and Environmental Studies, Kyoto University, Kyoto, Japan; Laboratory of Motor Control and Learning, Graduate School of Human and Environmental Studies, Kyoto University, Yoshida-nihonmatsu-cho, Sakyo-ku, Kyoto, Japan; Unit of Synergetic Studies for Space, Kyoto University, Kyoto, Japan

**Author notes:** Author Contributions Conceptualization TF, SH, MK Data Curation TF, SH, MK Formal Analysis TF, SH, MK Funding Acquisition MK Investigation TF, SH, MK Methodology TF, SH, MK Project Administration TF, SH, MK Resources SH, MK Software TF, SH Supervision TF, SH, MK Validation TF, SH, MK Visualization TF, SH, MK Writing – Original Draft Preparation TF Writing – Review & Editing TF, SH, MK.

## Abstract

Human bipedal posture is inherently unstable, making even daily activities potentially lead to falls and serious injuries. Although prior studies have shown that appropriate postural control supports both task success and postural balance during quiet standing or under modest postural demands, it remains unclear how the central nervous system controls whole-body posture under high-demand, near-fall conditions. Here, we investigated how varying postural demands influence postural strategies using a whole-body task in which participants leaned their body mediolaterally to reach a target. We manipulated the required leaning angles and velocities by varying target positions and time constraints to reach a target, thereby introducing different levels of postural demand. The results demonstrated that target position, time constraint, and movement distance significantly affected task performance, defined as reaching accuracy. Specifically, participants could accurately reach targets requiring upright or moderately leaning postures. However, when targets required greater leaning postures, participants failed to reach them. Furthermore, the detrimental effects of shorter time constraints and longer movement distances on task performance became more pronounced when target positions required greater leaning postures. These findings suggest that the central nervous system tolerates low to moderate postural demands to achieve task goals. In contrast, when postural demands exceed a certain threshold, the central nervous system begins to prioritize postural safety over task success. This study highlights the nonlinear effect of postural demands on motor planning during whole-body movements.

## Introduction

Maintaining standing postures is fundamental to our daily lives, as falls can lead to serious injury or worse. However, there are situations in which we intentionally take the risk of falling. For example, when a basketball player attempts to catch a ball heading out of bounds, the player has to lean significantly while keeping their feet inside the court. In such scenarios, the body’s center of mass (CoM) moves close to the stability limits generally defined by the area enclosed by the feet. However, to prevent falling, it is necessary to ensure that the CoM remains within the stability limits [1]. This posture thus carries a high risk of falling, as even a slight deviation can result in a fall. Although the central nervous system (CNS) is thought to consider the risk of falling when generating whole-body movements [2], the specific mechanism by which the CNS estimates and employs the risk is poorly understood. Therefore, how the CNS controls postural balance under near-falling conditions remains unclear.

Postural balance relies on effective CNS control [3, 4] in the presence of several destabilizing factors. For instance, sensorimotor noise [5, 6] or internal torque from the arm [7, 8] causes postural instability even when quiet standing or raising the arm in an upright posture. Additional task demands further complicate balance control, as greater trunk movement increases gravitational torque and the reduced stability limits constrain CoM movements [9, 10]. Previous studies have shown that postural control can adapt remarkably well to such challenging conditions [11-14]. For example, even under these greater postural demands, endpoint accuracy of reaching during standing did not change compared to lower postural demand conditions such as seated reaching [15]. These cited studies have generally focused on how the CNS compensates for destabilizing factors that cause deviation from a limited postural orientation, such as upright or moderately leaning postures in which the CoM did not approach the stability limit [16]. However, when higher postural demands, such as near-falling postures, are required, it remains unclear what postural control strategy the CNS would select. If the CNS considers the risk of falling when generating movements [2], it may prioritize fall avoidance by preventing the CoM from approaching the stability limits, rather than achieving the required near-falling posture accurately. To understand the postural control mechanism comprehensively, it is important to clarify whether and how the CNS changes its strategy in response to various conditions, including high demands such as a near-falling posture.

With this motivation, we constructed a novel task in which participants used their upper body to reach a target and were exposed to various postural demands. Postural demand was defined by the magnitude of the destabilizing torque. Considering the whole body as an inverted pendulum, the magnitude of the destabilizing torque is modulated by the position and acceleration of the CoM. Specifically, as the CoM position is close to the stability limits, the destabilizing torque increases due to gravity [9, 10]. Additionally, when the CoM acceleration increases, the destabilizing torque toward the stability limits also increases [9]. In this study, due to the nature of our experimental task, which required participants to dynamically control their posture while reaching toward a target using their upper body, we manipulated the position and acceleration of the center of the upper body (CoUB), rather than the CoM, to modulate postural demands. To systematically manipulate the positions and accelerations of the CoUB, we controlled the target positions and the time constraints to reach a target. For instance, when the target is positioned farther from an upright posture, the required CoUB position becomes closer to the stability limits (i.e., a near-falling posture), leading to high postural demand. On the other hand, since it was challenging to directly require an arbitrary CoUB acceleration, we indirectly modulated the CoUB acceleration by requiring higher CoUB velocity. When the time constraint to reach a target is shortened, a greater CoUB velocity is required to cover the same distance, consequently leading to greater CoUB acceleration. To summarize our approach, varying target positions and time constraints enabled us to systematically manipulate postural demands and compare postural control strategies in response to various postural demands including high-demand conditions such as a near-falling posture.

This study aimed to clarify postural control strategies in near-falling scenarios. We investigated how biomechanical postural demands affect postural control strategies by comparing the strategies in response to different target positions and time constraints to reach a target. Our study gives insights into understanding the control mechanisms during goal-directed whole-body movements.

## Materials and Methods

### Participants

Twenty healthy adults (all males, age: 23.2±1.4 years, height: 1.71±0.01 m, body mass: 62.3±5.9 kg, mean ± standard deviation [SD]) participated in this study. Participants approximately 1.70 m in height were recruited to control the effect of the body’s physical properties. Consequently, the mean height of participants was 1.71 m (range: 1.68 – 1.73). All participants had normal or corrected-to-normal vision and no history of musculoskeletal or neurological disorders. The participants provided written informed consent prior to the experiment. The recruitment period was from 29th October 2021 to 17th November 2021. This study was conducted by the Local Ethics Committee of the Graduate School of Human and Environmental Studies, Kyoto University (Approval Number 21-H-14).

### Experimental setup

Participants stood barefoot on a force plate (TF-4060, Tec Gihan Co. Ltd., Kyoto, Japan) with their feet positioned parallel and 30 cm apart, defined by the distance between the lateral edges of the feet. The foot position was marked to ensure a consistent stance throughout the experiment. The position of the center of foot pressure (CoP) was calculated based on the three components of the ground reaction force (*F_x_*, *F_y_*, and *F_z_*) and moment (*M_x_*, *M_y_*, and *M_z_*) that were recorded at 1000 Hz sampling frequency stored on a computer via a 16-bit A/D converter (NI-DAQ USB-6229, National Instruments, Austin, TX). Participants wore a head-mounted display (HMD; VIVE Pro 99HANW009-00, HTC, Tokyo, Japan) to interact with a virtual reality (VR) environment created using Unity (version 2019.3.9f1; Unity Technologies, California) (Fig 1A). The VR environment was used to eliminate task-irrelevant visual information. A screen (175 cm × 118 cm, 60 Hz refresh rate) was placed 2.3 m in front of participants in the VR environment.

**Fig 1.**
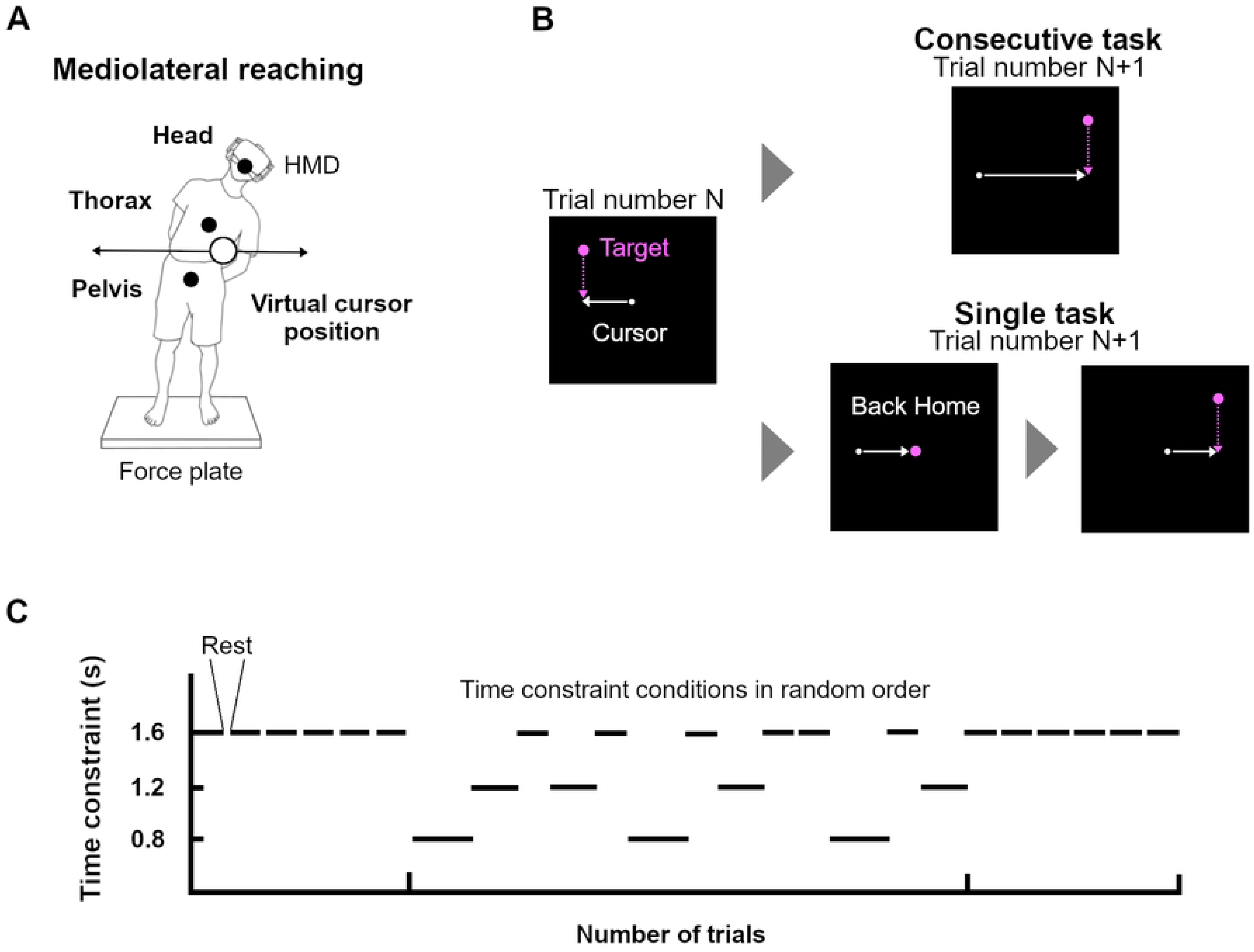
Experimental Setup. (A) Participants stood on a force plate and swayed mediolaterally while wearing a head-mounted display (HMD). The motion capture system acquired the mediolateral positions of the participant’s head, thorax, and pelvis to transform the mean position to the cursor position. (B) The screen was displayed through the HMD in front of the participants in virtual reality. Targets appeared at a fixed vertical height and moved downward. Participants swayed mediolaterally to move the cursor along a fixed horizontal path to reach the target. The mediolateral positions of the targets were at random and evenly distributed from side to side of the screen. In the consecutive task, the target appeared immediately after the previous one. In the single task, the target appeared at the center of the screen (home position), and the next target appeared once the cursor was within 1 cm of the home target. (C) Participants completed 2000 trials, including two single tasks and three consecutive tasks. After completing the single task for 400 trials (pre-single phase), participants performed the three consecutive tasks for a total of 1200 trials (consecutive phase) and then completed the single task for 400 trials again (post-single phase). Each single task included only the 1.6-second time constraint condition. The consecutive task consists of three time constraint conditions of 0.8, 1.2, and 1.6 seconds. Each condition had a total of 400 trials, divided into multiple blocks. The blocks in the consecutive task were presented in random order for each participant.

A cursor (diameter: 3.2 cm) displayed on the screen moved only in the mediolateral direction. The cursor position represented the spatial average across the mediolateral positions of six landmarks: (1) right earmuff of the HMD, (2) left earmuff of the HMD, (3) the xiphoid process, (4) the vertebral prominence of the seventh thoracic vertebra, (5) the center of both anterior superior iliac spines, (6) the sacrum. These landmarks, located symmetrically on opposite sides of the body axis, were selected to ensure cancellation of body rotational effects by averaging these positions. Thus, in our task, the cursor reflected mediolateral movements in the center position of the upper body across the head, thorax, and pelvis. Each body landmark was tracked with rigid bodies created by four or five infrared reflective markers. The marker positions were sampled at 100 Hz using a 3D optical motion capture system (OptiTrack V100, Natural Point Inc., Oregon, USA). The motion data were streamed in real-time from Motive 2.0.2 software (Natural Point Inc., Oregon, USA) via NatNet SDK to LabVIEW (National Instruments, Austin, TX), enabling real-time cursor calculation and display using the LabVIEW program.

### Whole body task

Participants performed a whole-body task to investigate how postural demands affect postural control strategies. Postural demand was defined by the magnitude of the destabilizing torque, which was modulated by two factors: (1) how close the position of the CoUB was to the stability limits that were assumed to be the area enclosed by the feet [9, 10, 17], (2) the acceleration of the CoUB [9, 10]. The task manipulated the two factors by varying target positions and the time constraints to reach a target. Before the task, participants were asked to stand upright on the force plate and the cursor position when participants stood upright was set at the center of the screen, referred to as the home position. Additionally, the line on which the cursor moved mediolaterally is referred to as the cursor movement line. In each trial during the task, a target (diameter: 9.6 cm) appeared at 22.4 cm above the cursor movement line and moved downward. The target disappeared when the target arrived at the cursor movement line. Participants were asked to control the cursor to reach the target while keeping their feet flat (Fig 1B left). To manipulate postural demand during the task, we varied the mediolateral target positions, which influenced the CoUB positions. For example, reaching a target far from the home position required participants to lean significantly laterally. Therefore, considering the whole body as an inverted pendulum [9, 10], the magnitude of the destabilizing torque increases as the CoUB position moves closer to the stability limits. Given that the home position was zero, the mediolateral target positions ranged from ±30.8 cm to the left and right, which was determined through a pilot study to include target positions unreachable without lifting a foot to induce near-falling situations. To cover the entire range, a total of 400 targets were used. The range was divided into 40 areas, and 10 target positions were randomly selected within each area. Consequently, target positions were presented in a random order within ±30.8 cm (Fig 2A). To manipulate postural demand, we also varied the time constraints to reach a target, which influenced the CoUB acceleration. The time constraint was defined as the time interval between the target’s appearance and disappearance. Therefore, the shorter time constraint required participants to lean with the greater CoUB velocity to cover the same distance, leading to greater CoUB acceleration.

**Fig 2.**
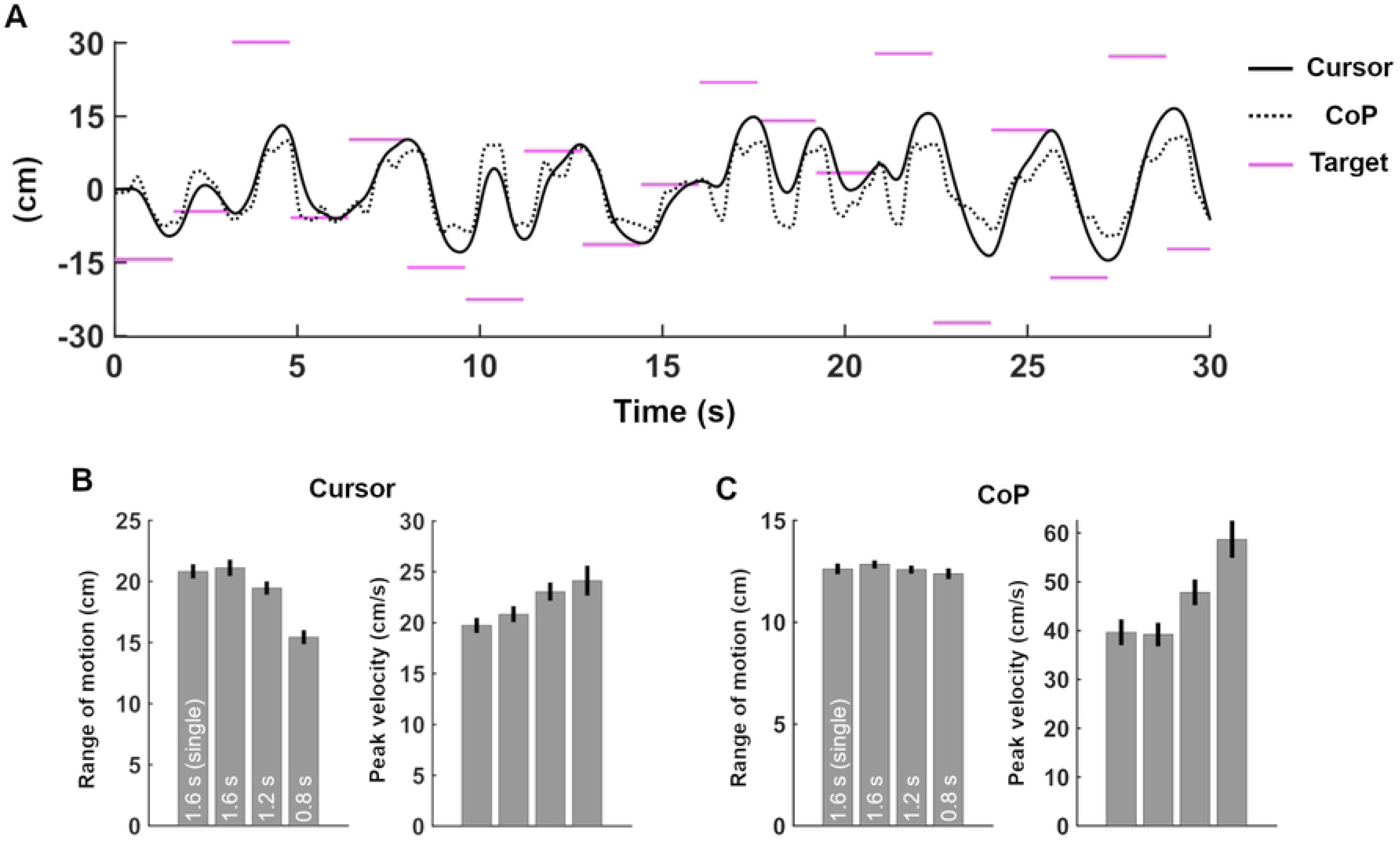
Trajectories and movement kinematic parameters. (A) Representative time series of cursor and CoP trajectories for the 1.6-second condition in the consecutive task. The solid black line indicates cursor movement; the dashed line indicates CoP movement. Each pink horizontal line represents a target position. (B) Group means of cursor range of motion and peak velocity across all conditions. Values are mean ± SEM across participants. Range of motion values were presented as absolute values, collapsed across left and right directions. (C) Group means of CoP range of motion and peak velocity across all conditions. Values are mean ± SEM across participants.

In the experiment, there were two task types: single task and consecutive task. In the single task, after each trial, the target returned to the home position before the next one appeared (Fig 1B bottom row). When the cursor was within 1 cm of the target at the home position, the next target appeared. When participants reached the target at the home position, there was no time limit so that participants could start the next trial at their own pace. On the other hand, in the consecutive task, after each trial, the next target immediately appeared and started to move (Fig 1B top row). The consecutive task was adopted to examine postural control strategies in a dynamic environment. Throughout the experiment, when the cursor was within 1 cm of the target, a pleasant chime sound was played from the earmuffs of the HMD (1400 Hz tone, lasting for 200 ms), and the target color changed from pink to blue to signify success. Participants were instructed to achieve as much success as possible and keep their feet flat on the force plate and their hands folded behind their back.

### Experimental Procedure

The experiment consisted of three phases: a pre-single phase, a consecutive phase, and a post-single phase. Before the pre-single phase, participants practiced controlling the cursor for 2-3 minutes, during which the participants could freely move the cursor. In the pre-single phase, participants completed a single task. A total of 400 targets were presented, and each trial’s time constraint was 1.6 seconds. There were 6 blocks every 75 trials for 2-3 minutes. The duration of each block depended on the participant’s pace, as there was no time limit when the target was in the home position in the single task.

In the consecutive phase, participants completed a consecutive task under three time constraint conditions (0.8, 1.2, and 1.6 seconds). 400 targets in each condition were presented. One block had different numbers of trials across time constraint conditions (1.6 s, 75 trials; 1.2 s, 100 trials; 0.8 s, 150 trials; Fig 1C). This ensured that each block lasted 2 minutes.

In the post-single phase, participants completed a single task again, allowing us to assess learning effects by comparing task performance between the pre-single phase and the post-single phase (see S1 Fig).

Participants took rest breaks for 1 minute between blocks and 3 minutes between phases. Additional rest time was allowed if the participants needed it. The experiment was interrupted if the participants’ feet left the force plate. If the experiment was interrupted, it was resumed after the feet were aligned with the marked positions. Data from interrupted trials were excluded from the analysis, resulting in the removal of 22 trials in all participants and phases.

### Data analysis and statistical analysis

The cursor data were low-pass filtered at 4 Hz using a zero-phase fourth-order Butterworth filter. The mediolateral CoP data were low-pass filtered at 10 Hz using a zero-phase fourth-order Butterworth filter. Each participant’s range of motion for the cursor and CoP position was quantified by the maximum lateral displacement in each condition. The cursor and CoP velocity were calculated using a three-point differentiation algorithm. Positive velocity values indicated movement toward the target. Concerning the cursor and CoP velocities, the peak velocity in each trial was quantified, and the mean peak velocity across trials was calculated for each participant and condition.

We quantified task performance based on an error between the cursor and the target positions when the target arrived at the cursor movement line. Positive values indicated undershooting in the direction of the home position, and negative values indicated overshooting in the direction of the stability limits. To investigate how the postural demands of the CoUB position within the stability limits affect errors, we compared errors in response to various target positions within each participant’s range of motion of the cursor. For each target area, we calculated the mean error for each participant. We combined left and right target positions relative to the home position, as direction did not affect the errors. A critical point was defined as the target position at which the lower limit of the 95% confidence interval of the mean exceeded zero for each participant. The group mean of errors and the 95% confidence interval of the group mean were calculated across all participants for each target area. Due to individual differences in the range of motion, data for the 16th and subsequent target areas were obtained from only four participants. To mitigate the influence of individual variability on the mean calculation, we reported the mean error only for target areas up to the 15th, where the number of participants exceeded half of the total (n ≥ 12).

To examine how the postural demands of the CoUB acceleration affect errors, we compared errors across different time constraint conditions. When comparing errors across conditions, we accounted for the effects of target positions and movement distances. Movement distance was defined by the distance between the target and the cursor when the target appeared. Target positions were classified into five groups, each group consisting of three adjacent target areas. This classification was based on the fact that more than half of the participants (n ≥ 12) had a range of motion that included up to the 15th target area (see Fig 2). Thus, we divided all target areas into five groups, each with three areas. The data after the 16th area was included in the 5th target position group. After classifying target positions, we then analyzed the effects of movement distance. To do so, we partitioned the distribution of movement distances into 5 bins for each condition and target group, ensuring that each bin contained approximately equal numbers of samples. Finally, we quantified the mean error for each bin to examine how movement distance influences errors. The above partition of target positions and movement distances was done for visualization purposes. Therefore, we performed statistical analysis on the effect of target positions and movement distances on errors across conditions as explained below.

To test the idea that target positions and time constraints affect errors, we fitted a generalized linear mixed-effects model. The dependent variable was an error. Three independent variables were target position, time constraint, and movement distance. The model equation was:

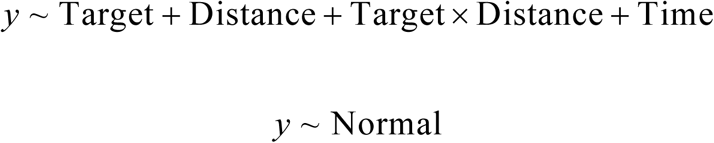

The model included participants as a random intercept. To minimize the influence of physical or physiological factors, such as joint range of motion or insufficient strength to move quickly, which could contribute to error, the data used in the model were limited to trials where the target positions fell within each participant’s range of motion of the cursor, and where the movement distance was less than the maximum distance covered by the cursor under each time constraint condition.

Data analysis was performed offline using MATLAB (R2024b, The MathWorks Inc., Natick, MA, United States). A generalized linear mixed-effects model was conducted using lme4 package [18] in R (version 4.2.2) and RStudio [19, 20]. In the statistical analysis, a *p*-value of less than 0.05 was considered statistically significant.

## Results

Fig 2A illustrates a representative time series of the cursor and CoP movement for the 1.6-second time constraint condition during the consecutive phase. The CoP movement generally coincides with the cursor movement. Fig 2B shows the group means of the range of motion and the peak velocity for the cursor movement for all conditions. As time constraints shortened, the range of motion was smaller (mean ± SE, single 1.6 s, 20.8 ± 0.60 cm; 1.6 s, 21.1 ± 0.68 cm; 1.2 s, 19.5 ± 0.56 cm; 0.8 s, 15.5 ± 0.59 cm) and the peak velocity increased (single 1.6 s, 19.8 ± 0.75 cm/s; 1.6 s, 20.8 ± 0.80 cm/s; 1.2 s, 23.1 ± 0.90 cm/s; 0.8 s, 24.1 ± 1.48 cm/s). Similarly, Fig 2C shows the group means of the range of motion and the peak velocity for the CoP movement for all conditions. While the range of motion was invariant for all conditions (mean ± SD, single 1.6 s, 12.6 ± 0.27 cm; 1.6 s, 12.8 ± 0.20 cm; 1.2 s, 12.6 ± 0.20 cm; 0.8 s, 12.4 ± 0.27 cm), the peak velocity increased as time constraints shortened (single 1.6 s, 39.6 ± 2.7 cm/s; 1.6 s, 39.2 ± 2.4 cm/s; 1.2 s, 47.8 ± 2.7 cm/s; 0.8 s, 58.7 ± 3.8 cm/s).

### Effect of target positions on errors in a single task

To clarify how target positions affect errors between the cursor and target positions, we compared errors for different target positions within each participant’s range of motion. Fig 3A shows errors in the single task during the post-single phase, where the time constraint was 1.6 seconds, and participants started reaching from an upright posture on all trials. We observed that errors increased as the target positions were lateral for all participants. Interestingly, despite being well before the limits of the range of motion, there was a critical point after which the errors increased sharply. Furthermore, Fig 3B demonstrates that the critical points varied by participants, although the participants had similar physical properties (mean ± SD, height: 1.71±0.01 m, body mass: 62.3±5.9 kg).

**Fig 3.**
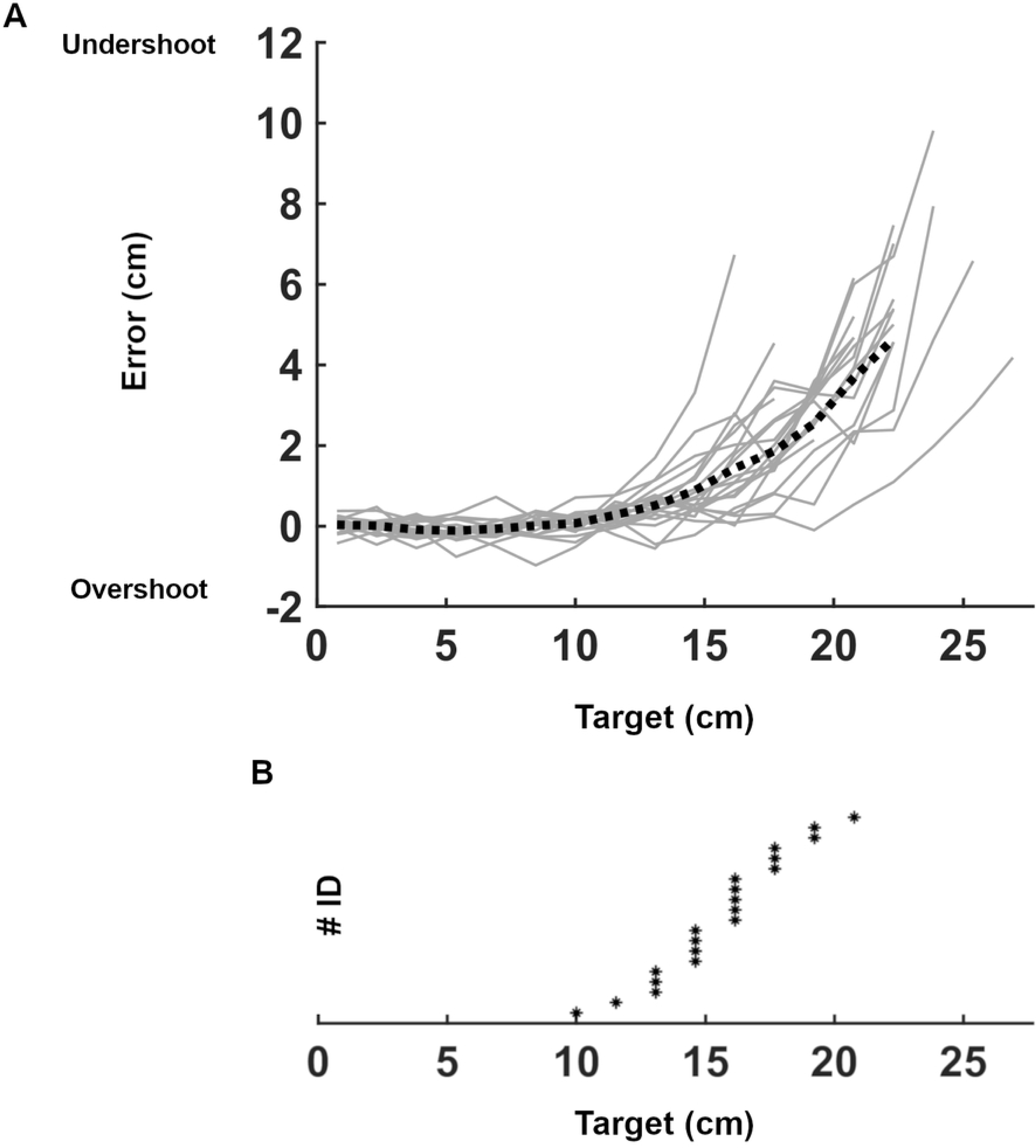
Task performance of the single task. (A) Error between the cursor and the target positions as a function of target position for each participant in the single task (post-single phase). Errors were calculated when the target reached the cursor movement line. Positive values indicate that the cursor was medial to the target (undershoot). Negative values indicate that the cursor was lateral to the target (overshoot). The dashed line represents the mean across all participants. Thin lines represent individual participants. (B) Asterisks denote the critical points at which the lower boundary of the 95% confidence interval exceeded zero for each participant.

In addition, S1 Fig shows the learning effect by comparing errors between the single tasks in the pre-single phase and the post-single phase. The more medial the target position, the more rapidly participants exhibited improvement in the first few trials of the pre-single phase. In contrast, the more lateral the target position, the slower the improvement was and continued throughout the experiment. Despite this slow learning effect, the increase in errors for lateral target positions observed in Fig 3A remained evident.

### Effect of target positions on errors in consecutive tasks

To examine how target positions affect errors for more dynamic postural control tasks, we compared errors between the 1.6-second condition of the consecutive task and the 1.6-second condition of the single task during the post-single phase. Fig 4A shows that the trend of errors was similar for two tasks. Moreover, Fig 4B demonstrates similar critical points between two tasks (mean ± SD, single 1.6 s, 15.6 ± 2.7 cm; consecutive 1.6 s, 15.7 ± 3.1 cm). In the single task, the initial postures were identical on all trials. Conversely, in the consecutive task, the initial postures varied depending on the previous trial. Thus, if target positions were the same, the initial postures differed between the two tasks, resulting in variations in movement distances from the initial cursor position to a target position. Despite these differences, we found the identical critical points between these tasks when the time constraint was 1.6 seconds, indicating the effect of target positions on errors (Fig 4B).

**Fig 4.**
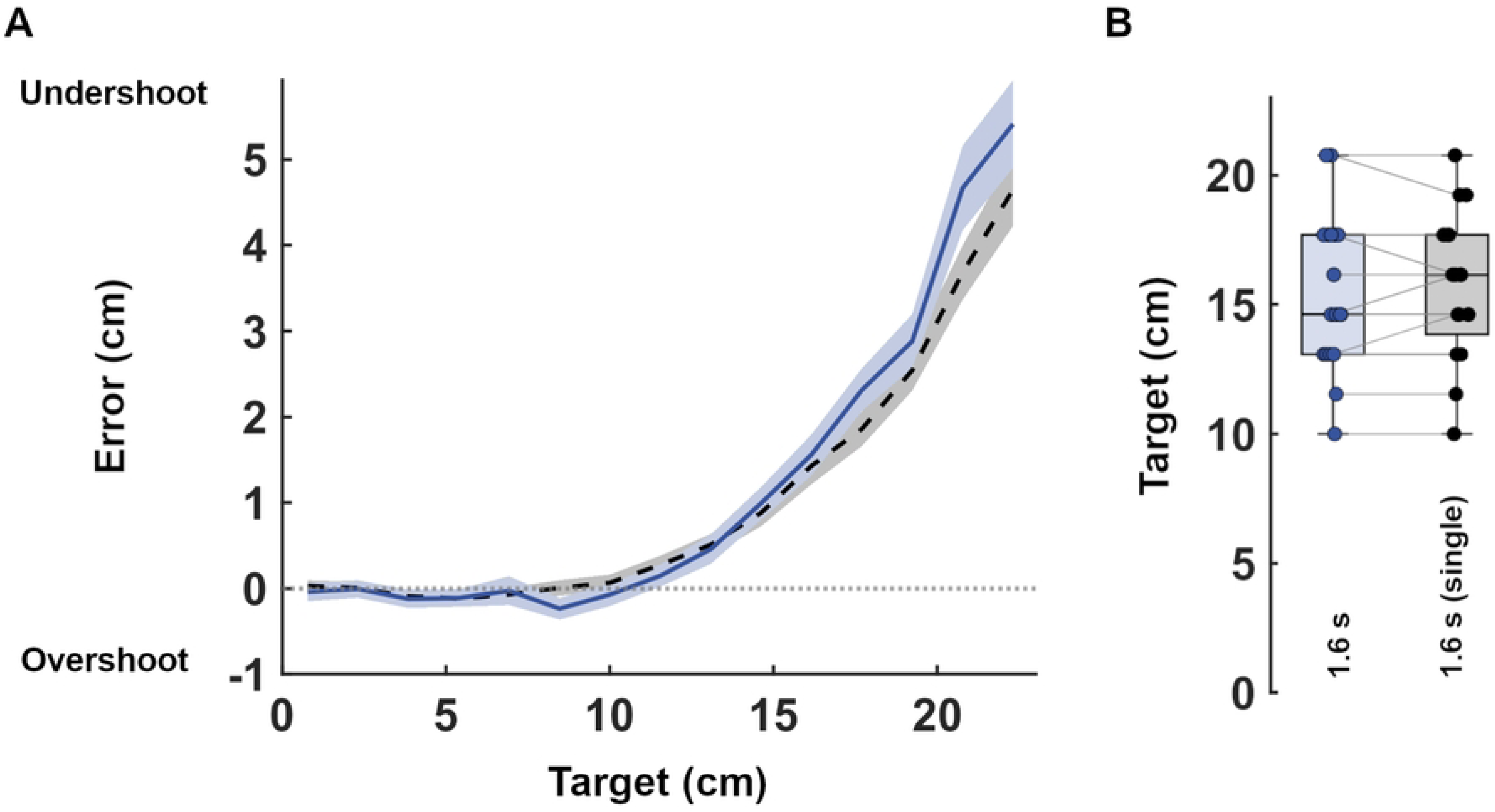
Comparison between the single task and the consecutive task. (A) Error as a function of target position. Lines and shaded areas represent the mean and 95% confidence interval across participants. The solid black line indicates the consecutive task (1.6-second time constraint condition), while the dashed line indicates the single task in the post-single phase. Data are shown only for target positions within the cursor range of motion for more than half of the participants (n ≥ 12). (B) Comparison of critical points. In the box plot, the midline, box size, and whiskers represent the median, 25th-75th percentiles, and the range within 1.5 times the interquartile range (IQR), respectively. Dots represent individual participants.

### Effect of time constraints on errors

Next, we examined whether and how time constraints affected errors. S2 Fig shows that, for all conditions, errors increased as target positions were lateral. However, as time constraints shortened, the critical point shifted toward the medial side (mean ± SD, 1.6 s, 15.7 ± 3.1 cm; 1.2 s, 12.1 ± 2.7, 0.8 s, 5.7 ± 1.9 cm), indicating that factors other than target positions affect errors. Thus, to examine the effect of time constraints on errors, we compared errors across different time constraint conditions by considering the effects of target positions and movement distances on errors. Fig 5 illustrates the relationship between errors and movement distances, presented separately for the five target area groups. First, as found in Fig 4, errors increased as target areas were lateral for all conditions. Second, errors increased as time constraints shortened for each target area group. Finally, for each target area group, errors increased as movement distance was greater, except for the most medial target area (Fig 5 most left).

**Fig 5.**
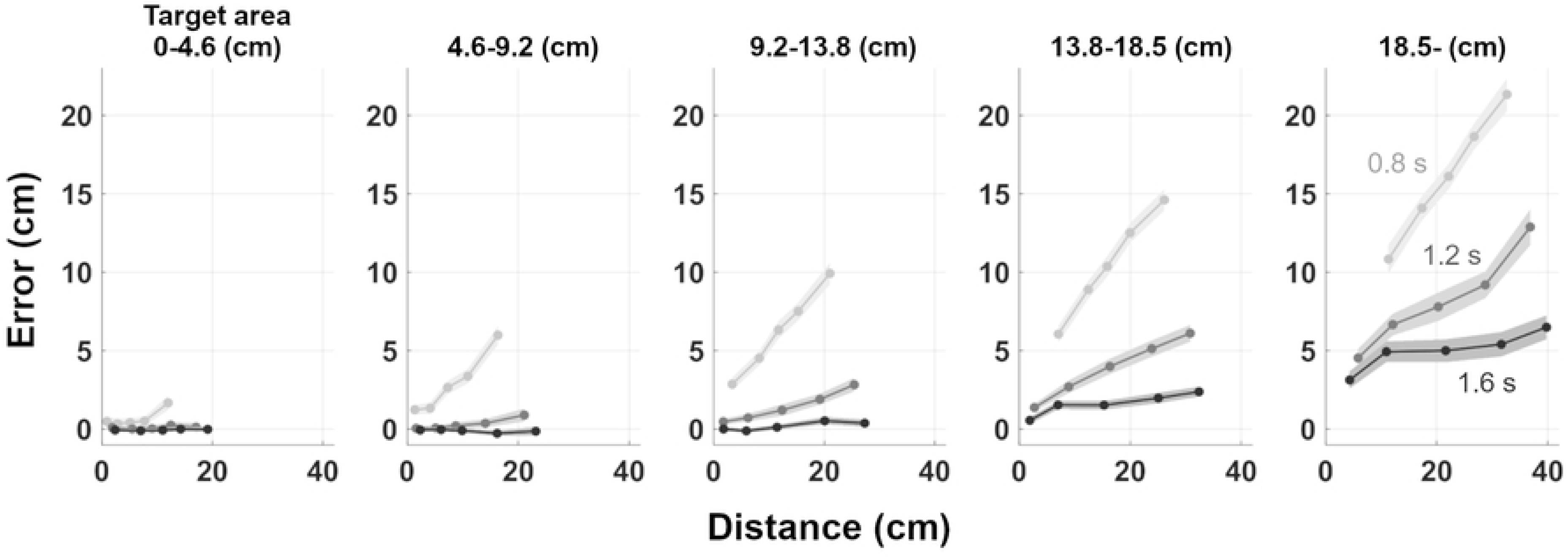
Relationship between distance and error for target areas and time constraint conditions. Error as a function of the distance between the target and cursor when the target appeared. Target positions were divided into 5 groups, each shown separately. Line darkness corresponds to different time constraint conditions. Shaded areas represent the 95% confidence interval of the mean. Data points are plotted at the center of distance bins, each adjusted to include approximately equal numbers of samples.

To test the idea that target position, time constraint, and movement distance affect errors, we performed a generalized linear mixed-effects model. Significant main effects were observed in target position (β = 0.26, SE = 0.0077, 95% CI = [0.24, 0.27], *p* < 0.001), time constraint (β = -0.073, SE = 0.00085, 95% CI = [-0.074, -0.071], *p* < 0.001), and movement distance (β = 0.049, SE = 0.0077, 95% CI = [0.033, 0.064], *p* < 0.001). In addition, the interaction between target position and movement distance was significant (β = 0.0082, SE = 0.00048, 95% CI = [0.0072, 0.0092], *p* < 0.001).

Collectively, the three factors—target position, time constraint, and movement distance—influence errors in our task. It is important to note that target position and time constraint affect errors significantly, even when considering movement distance due to different initial postures in the consecutive task.

## Discussion

The purpose of this study was to identify postural control strategies in response to varying biomechanical postural demands, including near-falling situations. We manipulated postural demand by requiring various positions and accelerations of the center of the upper body using different target positions and time constraints to reach a target. During our experiment, some participants experienced falls defined based on whether the participants left their foot from the force plate to reach a target. This indicates that our task encouraged the participants to be in near-falling situations. In addition, we observed that the peak cursor velocity decreased as the time constraints shortened (Fig 2B), indicating that our task manipulated the postural demands by varying time constraints to reach a target. We quantified errors between the cursor and target positions as task performance. Our main finding was that target positions and time constraints significantly affect errors. Additionally, we observed a large individual difference in the critical point of target positions at which errors began to increase sharply. We will discuss below why errors increased as target positions became more lateral and time constraints shortened. These findings suggest that the CNS adopts a conservative strategy to account for the risk of falling in response to the biomechanical postural demands, that is, the destabilizing torque applied to the body.

### The risk of falling based on CoM positions

We observed significantly large errors beyond the critical point (Figs 3 and 4). This indicates that the CNS prioritized postural objectives to maintain the body as upright as possible, over task success. This supports the earlier suggestion that the CNS considers the risk of injury, such as falls, in generating movements [2]. Babič et al. found that, in a motor adaptation task, the residual difference between the original and adapted trajectories was significantly larger in a whole-body task than in a seated arm-reaching task [2]. They concluded that this difference was due to the CNS accounting for the destabilizing effect of gravity. Our study can give further insight into this suggestion. In our task, although gravity destabilized the various postures, errors increased only beyond the critical point (Figs 3 and 4). Thus, we suggest that the CNS prioritizes postural objectives to maintain the body as upright as possible only when the CoM position exceeds a certain threshold within the stability limits.

On the other hand, we observed that endpoint accuracy did not change across the target positions within a critical point (Figs 3 and 4). This result is also broadly consistent with previous works demonstrating a high adaptability of postural control against postural constraints. Several studies have manipulated postural constraints using approaches such as reducing the size of stability limits, requiring reaching beyond one’s reach, and/or introducing an unstable support surface. These studies reported that endpoint accuracy [11-15, 21] and muscle coordination [22, 23] remained unaltered compared to less challenging conditions. This high adaptability against postural constraints may align with the consistent endpoint accuracy within the critical point observed in our study. Indeed, the constraints in previous studies were designed to prevent the CoP from approaching the stability limits [16], resulting in a narrower range of CoP movement than that observed in our task (Fig 2C). Thus, the CoM positions required in previous studies were likely below the threshold. Taken together, previous findings and our results suggest that when the CoM positions remain below a certain threshold, the CNS does not consider the risk of falling in generating movements.

Moreover, we found a large individual difference in the critical points (Fig 3) despite our participants having similar physical properties (height: 1.71±0.01 m, body mass: 62.3±5.9 kg, mean ± SD). This finding suggests that the individual difference in the critical points is shaped by the CNS control process rather than being determined only by physiological factors. Such individual differences in motor strategies have been explained in various ways. For instance, one study explained the differences in terms of variations in internal values [24]. Other studies have described them as differences in risk sensitivity [25-29]. Additionally, studies on postural threat have explained differences in postural strategies due to fear of falling [30, 31]. Although it is challenging to identify the specific causes of the individual differences in our study, this finding supports our suggestion that the CNS estimates the risk of falling based on CoM positions to adopt a conservative strategy that prioritizes postural objectives to maintain the body as upright as possible.

### The risk of falling based on CoM velocities

In our study, we also examined the effect of postural demands regarding accelerations by comparing errors across time constraint conditions. Our result shows that the errors increased significantly as time constraints shortened, even if the target positions and movement distances were identical across the conditions (Fig 5). This result indicates that the CNS did not generate the necessary movement velocity to reach the target. We suggest that the CNS might prefer to suppress the CoM velocity at the expense of task success, by estimating the risk of falling from the CoM velocity information. Computational studies support this hypothesis, showing that quiet standing measured experimentally can be explained well by a postural control system using the CoM velocity information as a feedback variable [5, 6, 32]. By estimating fall risk from CoM velocity, the CNS may adopt a conservative strategy to maintain stability in dynamic environments.

However, it is possible that an alternative explanation exists. Previous studies found that unidirectional CoP and CoM movements had a mean duration of around 1 second during quiet standing [33, 34]. It is suggested that this reflects the characteristics of intermittent control by the CNS since the mean duration did not alter in response to different dynamics requirements [35, 36]. On the contrary, in our study, the mean duration of CoP movements modulated in response to different time constraint conditions (0.8 s condition: 0.40 s, 1.2 s condition: 0.45 s, 1.6 s condition: 0.48 s, see S3 Fig). Therefore, the apparent limit of movement velocity may occur because the CNS considers the risk of falling based on the CoM velocity.

## Conclusion

In conclusion, this study examined postural control strategies in response to various postural demands. The results demonstrated that target positions, time constraints, and the movement distances influenced the task performance, quantified as the positional error between the cursor and the target. The error increased sharply when the required target positions exceeded an individual-specific threshold. Moreover, the error increased as the time constraints shortened and the movement distances increased, especially when the target positions were located more laterally relative to the upright posture. These nonlinear effects suggest that the CNS begins prioritizing postural objectives to maintain the body as upright as possible over task success, only when the postural demands surpass a critical threshold. Such insights could have important implications for understanding the control mechanisms during goal-directed whole-body movements.

## Supporting information

S1 Fig. Learning effect.

S2 Fig. Task performance for time constraint conditions.

